# Speech, gait, and brain dynamics during natural conversation in motion

**DOI:** 10.64898/2025.12.23.696153

**Authors:** Martin Orf, Sarah Tune, Ronny Hannemann, Jonas Obleser

**Affiliations:** Department of Psychology, University of Lübeck, Lübeck, Germany; Center of Brain, Behavior and Metabolism (CBBM), University of Lübeck, Lübeck, Germany; ORCA Labs Europe, Erlangen

**Keywords:** Neural Speech tracking, EEG, Walking, Gait, Speech production and perception, Real-world

## Abstract

Communication connects us to others and immerses us in our environment. Mobile EEG and hearing aid-embedded motion sensors allow investigation of communi-cation and movement in real-world contexts. Here we reliably record and model neural responses to natural conversational speech while participants are walking, demonstrating the feasibility of combining EEG with whole-body movement moni-toring. Participants (N = 61) performed focused listening and natural conversation while walking indoors and outdoors. Motion sensors, EEG, and audio recordings captured behavioural, neural, and motor dynamics. Walking speed varied with en-vironment and task: faster outdoors and slower during speaking. Temporal align-ment between gait and speech revealed coordinated rhythmic interactions, stronger during speaking than listening. Auditory temporal response functions were preserved across listening tasks. Gait TRFs displayed lateralized and temporal dy-namics consistent across tasks. These results provide insight into the coordinated interplay of speech, brain activity, and movement. This highlights the potential of multimodal approaches in ecologically valid settings.

## Introduction

Engaging in conversation while walking through dynamic environments—such as bustling city streets or crowded public spaces—is a common aspect of daily life. These situations demand the simultaneous coordination of motor and cognitive processes, as individuals navigate their surroundings and maintain social interac-tions (Al-Yahya et al., 2011). Understanding how the brain supports such complex, multimodal tasks is crucial for advancing our knowledge of human cognition and developing technologies that assist individuals in real-world settings (Jungnickel & Gramann, 2016).

Natural conversation often unfolds in dynamic, multimodal contexts where speech and movement occur simultaneously. During such interactions, the rhythm and temporal structure of speech, as captured by the speech envelope, may interact with ongoing gait patterns. Speakers might adjust their walking pace to match their own speech rhythm, or listeners could subtly align their gait to the envelope of the conversational partner, facilitating coordination and turn-taking. This bidirectional coupling between speech and locomotion could reflect an embodied strategy to optimize communication in real-world settings. However, empirical studies directly linking speech envelope dynamics to gait patterns during natural conversation re-main scarce.

Building on this behavioural perspective, recent advances suggest that brain ac-tivity, as measured with electroencephalography (EEG), can be leveraged to sup-port auditory processing in complex listening situations. Specifically, low-frequency features of speech can be tracked in the brain—a phenomenon referred to as neu-ral speech tracking—and are modulated by selective attention (Ding & Simon, 2012; Obleser & Kayser, 2019; O’Sullivan et al., 2015). By evaluating the correla-tion between EEG signals and speech envelopes, it is possible to identify which speaker a listener is attending to, enabling brain-guided auditory enhancement (Orf et al., 2023; Tune et al., 2021; Tune & Obleser, 2024; Zion Golumbic et al., 2013). However, most studies have been conducted in stationary, highly controlled labor-atory environments, where neither listeners nor sound sources move, limiting eco-logical validity. To better understand how the brain supports real-world communi-cation, hearing science and cognitive neuroscience must move beyond the labor-atory, as everyday listening occurs in dynamic, multimodal, and often outdoor en-vironments (Hamilton & Huth, 2020). Recently, this approach has increasingly been implemented, and investigating brain mechanisms during natural movement and real-world activities has gained considerable relevance in neuroscience (Bigand et al., 2025; Haupt et al., 2025; Levy et al., 2025; Schmidt-Kassow et al., 2025; Wöstmann & Obleser, 2025).

Beyond auditory processing, real-world listening often involves simultaneous mo-tor activity such as walking, turning, or gesturing, which can act as a secondary task and increase cognitive load, potentially impacting attentive listening (Al-Yahya et al., 2011; McPhee et al., 2022). Ensuring reliable EEG measurements in such conditions requires careful selection of hardware and analysis strategies, as em-phasized by Gramann et al. (2024), to maintain data quality and validity outside laboratory settings. Despite challenges such as motion-induced EEG artifacts (Ja-cobsen et al., 2021), mobile EEG systems have been shown to reliably capture neural correlates of cognitive and auditory processing in freely walking individuals (De Vos et al., 2014; Debener et al., 2012; Reiser et al., 2021). Recent work by Straetmans et al. (2024) has shown that EEG-based auditory attention decoding remains robust even in complex real-world environments such as busy cafeterias and urban streets , using listening-to-audiobook paradigms. However, whether similarly reliable neural responses can be observed during genuine conversational interactions, which involve dynamic alternations between listening and speaking, remains an open question. In the present study, we directly address this gap by investigating neural responses during natural conversation, encompassing both listening and speaking.

While auditory neural responses during complex real-world listening have begun to receive increased attention, research on gait-related ERPs remains scarce. Ex-isting studies, such as Herbert & Munz, (2020), have primarily examined gait-re-lated neural activity under highly controlled laboratory conditions, for example dur-ing treadmill walking. As a result, little is currently known about how gait and com-munication interact, or about the neural correlates of this interaction in ecologically valid, naturalistic settings.

Here, we investigated a potential link between the envelope of self-produced speech and own gait, between the speech rhythm of the conversational partner and own gait. We hypothesized a coupling between a speaker’s gait and their pro-duced speech, and likewise a coupling of a listener’s gait to the speech envelope of their partner. Specifically, we expected that gait patterns would align to the tem-poral dynamics of speech. Furthermore, we were interested in the neural repre-sentation of self-produced and externally produced and gait occurring simultaneously occurring from a natural conversation. We asked whether reliable temporal response functions (TRFs) could be obtained during natural conversation for both speaking and listening, and whether TRFs could also be observed for par-ticipants’ gait.

To address these questions, we conducted a semi-naturalistic dyadic conversation experiment in which participants walked while engaging in a semi-scripted dialogue with an experimenter. Participants wore a hearing aid (amplification off) and a port-able EEG system, both equipped with motion sensors, allowing synchronized re-cording of gait, EEG, and audio. Conversations were recorded with microphones on both participant and experimenter. In addition to the conversation, participants completed control tasks—Focused Listening, a Digit Triplet Test, plus a walking-only Baseline—performed indoors and outdoors. For the present study, we focus on the Conversation and Focused Listening tasks to investigate speech–gait inter-action and its neural correlates under ecologically valid conditions.

## Methods

### Participants

Sixty-one adults (30 female, 31 male) aged between 40 and 80 years (mean age: 65.7 years) participated in the study. All participants were native speakers of Ger-man and reported normal hearing to mild or moderate age-related hearing loss (presbycusis), with no history of neurological disorders. Pure-tone audiometry was conducted at frequencies between 125 and 8000 Hz to verify hearing status. Across the tested frequencies, all participants showed auditory thresholds below 40 dB HL (mean: 19.3 dB HL). To assess physical fitness, participants completed the Short Physical Performance Battery (SPPB), which includes a balance test, a gait speed test, and the timed up-and -go test. All participants successfully passed the assessment. Data from fifteen participants were excluded from analyses rely-ing on microphone recordings due to corrupted audio files. As a result, neural anal-yses of the focused listening task included data from 46 participants. For the natu-ral listening task, four additional participants were excluded due to the absence of any detectable TRF modulation, resulting in a final sample of 42 participants for this condition. All participants provided written informed consent and received com-pensation of €12 per hour. The study was approved by the local ethics committee of the University of Lübeck.

### Experimental Procedure

The experiment comprised four walking-based conditions: conversation, focused listening, digit triplet test (DTT), and baseline walking. All conditions were carried out both indoors (hallway) and outdoors (courtyard).

In the conversation task, participants engaged in a semi-scripted dialogue with an experimental leader while walking. The script was structured to avoid participants sharing sensitive personal information and to ensure a balanced speaking ratio between experimenter and participant but otherwise allowed for free conversation. Each session lasted 20 minutes per environment.

In the focused listening task, participants listened to a professionally narrated au-diobook *Ludwig van Beethoven: Basiswissen*. Short repeated segments (400 ms) were included in the audiobook material (Orf et al., 2023, 2025). They were in-structed to indicate the repetitions with a button press (numpad). The task duration was 10 minutes. The experiment was created using Psychophysics Toolbox ex-tensions (Brainard, 1997) and MATLAB (MathWorks, Natick, MA, USA).

In the DTT, participants heard 25 sequences of three spoken digits (1-9) presented in varying levels of background noise and were asked to recall them. On average, this task lasted 8 minutes. The experiment was created using Psychophysics Toolbox extensions (Brainard, 1997) and MATLAB (MathWorks, Natick, MA, USA). All audio files were sampled at 44.1 kHz with 16-bit resolution.

In the baseline condition, participants walked without any additional task. They walked three 35-m segments, turning around after each segment. Importantly, the same walking pattern was applied in all other experimental tasks, both indoors and outdoors. The order of experimental conditions and the indoor/outdoor environ-ment was randomized and balanced across participants.

### Experimental Setup

The experiment was conducted under two environmental conditions: inside (a hall-way of a research building) and outside (a courtyard of the same building). Partic-ipants wore the same technical setup across all environmental conditions and ex-perimental tasks (baseline, digit triplet task, focused listening, and conversation; Figure 1). They were equipped with a hearing aid (Signia HA Pure 312 7 NX GB; without amplification) containing motion sensors (accelerometer and gyroscope). In addition, they wore a mobile EEG cap (32-channel SMARTING Pro), which was also equipped with motion sensors (accelerometer and gyroscope). Participants also wore a neck loudspeaker (Panasonic; Model No.: SC-GN01) and a micro-phone (RODE Microphones Wireless GO II) for audio recording. The experimenter was additionally equipped with a microphone (RODE Microphones Wireless GO II). All participants carried a backpack containing a laptop (Lenovo; Version 23H2), which served both as the stimulus presentation and recording device. Data from all devices were synchronized via the Smarting recording software (PRO_072, mBrainTrain, Belgrade, Serbia), with the exception of the hearing aid motion sen-sors, which were later aligned using the motion data from the EEG system. Hearing aid data were recorded via app (Gait Study APP; Version 1.0) on a smartphone (Galaxy S20 FE, Modell No.: SM-G780G/DS). Due to the complexity of the setup, including the number of devices and parallel data streams, a temporal lag was introduced in the microphone recordings, which was identified using wired versus wireless microphone measurements and subsequently accounted for in the neural analysis using a re-alignment procedure (Carta et al., 2023).

**Figure 1.**
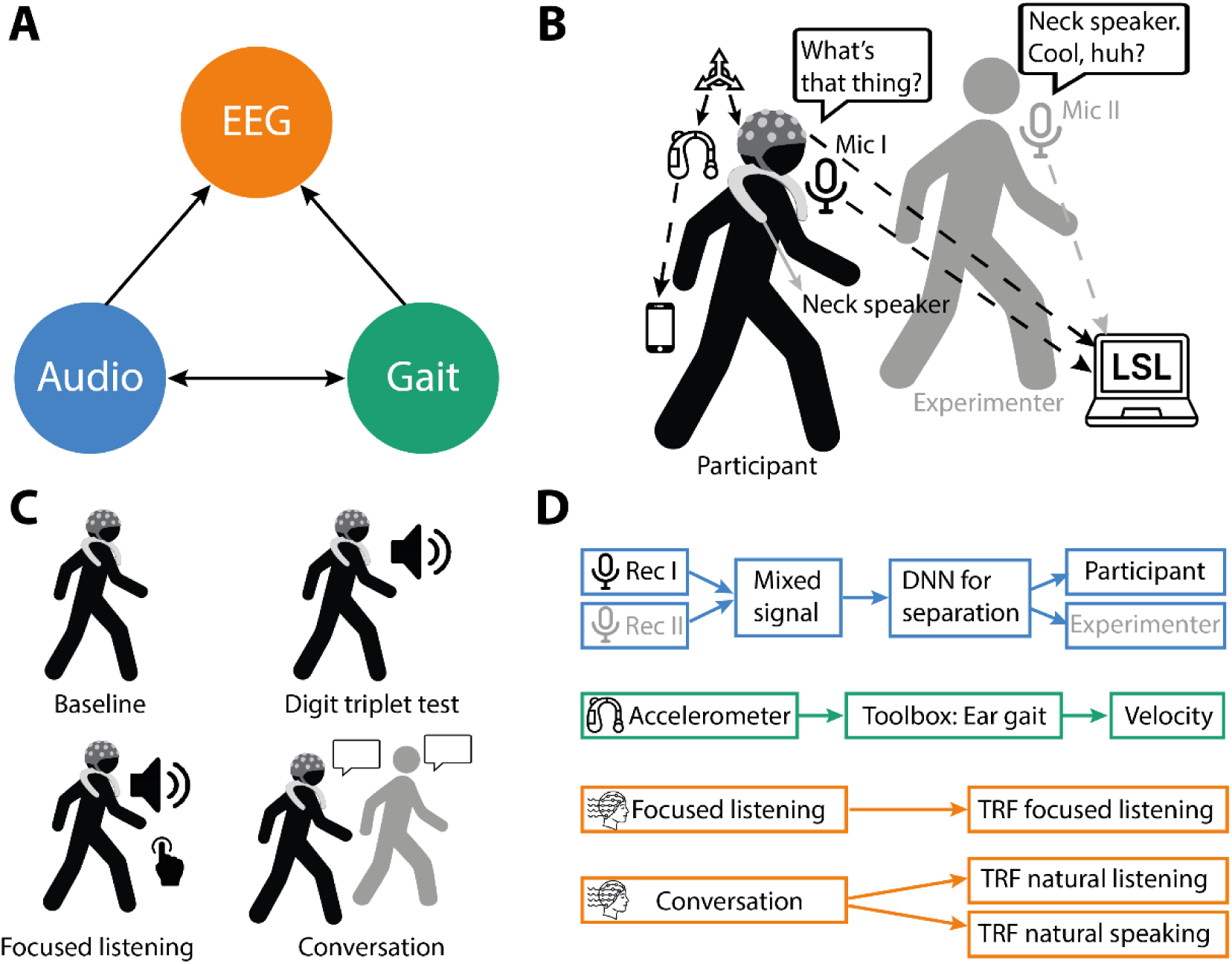
Experimental design and analysis framework. (A) Conceptual overview illustrating the triad of the three main modalities: EEG (orange), audio (blue), and gait (green). (B) Experimental setup: Participants wore EEG, hearing aid (motion sensors) and a neck-mounted speaker while walking. Audio was recorded from two microphones (participant and experimenter). Signals were streamed to the Lab Streaming Layer (LSL) for synchronization.(C) Experimental conditions (all during walking) included baseline walking, a digit triplet test (speech-in-noise), focused listening to an audiobook, and natural conversation with the experimenter.(D) Data processing pipeline: Audio recordings from participant and experimenter were separated using a deep neural network (DNN). Gait velocity was estimated from accelerometer data using the Ear gait toolbox. EEG data were analysed with temporal response functions (TRFs) to assess neural tracking during focused listening, natural listening, and natural speaking.

### Data acquisition and pre-processing Motion data

For gait-recording only, the hearing aid was positioned at the right ear for all par-ticipants, and the hearing-aid receiver was individually selected to ensure the best fit with respect to pinna size. Motion data from the embedded accelerometer and gyroscope were sampled at 50 Hz and further processed using the EarGait toolbox (Seifer et al., 2023, 2025). The preprocessing pipeline included alignment of the inertial data to the body-frame coordinate system, extraction of walking intervals, detection of gait events (initial contact, toe-off), and computation of temporal and spatial gait parameters such as gait velocity. For the continuous representation of gait, we used the pre-processed data from the superior–inferior (SI) axis, which also serves as the basis for gait parameter estimation in the EarGait toolbox.

### EEG data

A 32 electrode EEG-cap (Easycap, Herrsching, Germany; Ag-AgCl electrodes po-sitioned in accordance with the 10-20 International System) connected to a SMARTING amp was used to record the EEG (mBrainTrain, Belgrade, Serbia). This portable EEG system sends the signal via Bluetooth to a computer for record-ing. Using the program Smarting Streamer (mBrainTrain, version 3.4.2), EEG ac-tivity was captured at a sample rate of 250 Hz. Impedances were kept under 20 k**Ω** while impedances were used as an online reference during recording using electrode FCz. The Fieldtrip-toolbox, built-in functions, and MATLAB (Version 2018a Mathworks Inc., Natick, MA, United States) were used for offline EEG pre-processing (Oostenveld et al., 2011). High- and low-pass filters were applied to the EEG data between 1 and 100 Hz, and the electrodes M1 and M2 (the left and right mastoids) were averaged (two-pass Hamming window, FIR). On the EEG data from every participant, an independent component analysis (ICA) was performed.

Prior to ICA, M1 and M2 were removed. Only ICA artifacts related to eye blinks and eye movements were removed. Back projected to the data were elements not connected to artifacts. Clean EEG data were processed further. Frequencies up to 8 Hz are associated with neural speech tracking (Luo & Poeppel, 2007). EEG data were therefore low-pass filtered once more at 10 Hz (two-pass Hamming window, FIR). EEG data were then resampled to 125 Hz.

### Spoken audio recordings

Microphones (RODE Wireless GO II) for audio capture recorded at a sampling rate of 16 kHz. Although recorded under the same clock, the signals were captured as separate streams in the EEG recording software. We combined the recordings from both microphones (one worn by the participant, one by the experimenter) and then applied a deep neural network algorithm SpeechBrain (Ravanelli et al., 2021) to separate the two speakers. The resulting separated streams were used as the basis for computing the individual speech envelopes.

By calculating the onset envelope of each audio stream, the temporal fluctuations of speech were quantified (Fiedler et al., 2017). In the beginning, we used the NSL toolbox (Chi et al., 2005) to compute an auditory spectrogram (128 sub-band en-velopes logarithmically spaced between 90 and 4000 Hz).

To create a broad temporal envelope, the auditory spectrogram was secondly av-eraged across frequencies. Third, the half-wave rectified first derivative of the on-set envelope was obtained by computing the first derivative of this envelope and zeroing negative values. In order to match the EEG analysis’s target sampling rate, the onset envelope was lastly down sampled (125 Hz). For the speech gait rela-tionship, the onset envelops were down sampled to 50 Hz.

### Temporal response function

A temporal response function (TRF) is a linear filter model that describes how an input signal, such as the acoustic speech envelope, is transformed into a corre-sponding output signal, such as recorded EEG or continuous gait. To calculate the TRFs, we employed a multiple linear regression method (Crosse et al., 2016).

We applied the encoding TRF framework (forward model) in two analyses: the speech–gait coupling and the EEG analysis. For the speech–gait coupling, the separated speech envelopes served as input, and the continuous gait signal ex-tracted from the EarGait toolbox served as output. In this analysis, we examined temporal lags between changes in the speech envelope and the gait signal in a range from –1000 to +1000 ms. For the EEG analysis, the inputs were the sepa-rated speech envelopes and the continuous gait signal, while the output was the EEG recording. Here, we examined temporal lags between changes in the input signals and the EEG response in a range from –200 to +500 ms.

To prevent overfitting, ridge regression was used to estimate the TRFs. The opti-mal ridge parameter was determined separately for each participant using cross-validation. A predefined range of ridge values (λ = 10⁻⁶ to 10⁶ in logarithmic steps; Crosse et al., 2016) was tested, and a separate model was trained for each value using a 5-fold cross-validation procedure (split = 5). For each fold, the model was trained on four subsets and evaluated on the remaining subset by predicting the neural response, with predictions averaged across trials. The ridge parameter yielding the lowest mean squared error (MSE) across folds was selected as the optimal, subject-specific value.

To account for the temporal imprecision of our technical setup, we followed the re-alignment procedure by Carta et al. (2023). The procedure realigns data by fitting a TRF and shifting the data so that the strongest negative TRF component (as-sumed to be the N1) occurs at a fixed latency of 100 ms. This approach relies on the assumption that a clear N1 is present and inflates the N1 component by con-struction; therefore, results should be interpreted in terms of overall TRF shape, topography, and prediction accuracy rather than absolute latencies or N1 ampli-tude.

Prediction correlation refers to the Pearson correlation coefficient (r) between the predicted and measured gait signals and was used to quantify speech–gait cou-pling, using a leave-one-out cross-validation procedure. Encoding accuracy, in contrast, reflects how well a single stimulus stream (speech or gait) is represented in the EEG signal and was quantified using TRFs to predict the EEG response. Encoding accuracy was calculated as the Pearson correlation between the pre-dicted and the recorded EEG signals, also using a leave-one-out cross-validation procedure to predict the EEG (Crosse et al., 2016).

Importantly, encoding accuracy was computed using the uncorrected (non–rea-ligned) neural data, as it integrates information across a wide range of time lags and is therefore not expected to be affected by timing imprecisions in the experi-mental setup.

### Statistical analysis

We conducted statistical analyses on participants’ walking behaviour, the coupling between speech and gait, and their neural responses across conditions involving listening, speaking, and simultaneous walking, using linear mixed-effects models. These models account for the hierarchical structure of multi-condition, within-subject designs by incorporating both fixed and random effects. Predictors were centered and scaled where necessary, and random intercepts were included when appropriate to account for inter-individual differences.

Participants walking behaviour were analyzed in Matlab using the fitlme function and the following model:

***Gait velocity ∼1 +condition + Location + AGE + PTA + condition*Location + (1|subject)***

In this model, gait velocity served as the dependent variable, representing participants’ walking speed in m/s. *Condition* referred to the different task contexts (baseline, focused listening, DTT, and conversation), while *Location* indicated the environmental setting (indoors vs. outdoors). Participants’ age (AGE) and hearing loss (PTA) were included as covariates, with PTA defined as the average pure-tone threshold across 500, 1000, 2000, and 4000 Hz, averaged across the left and right ear.

The coupling between speech and gait was analyzed in Matlab using the fitlme function and the following models:

***EncAcc ∼ Condition * Location + Age + PTA + (1 | Subject)***

For listening and produced speech gait coupling, Condition varied accordingly (NonShifted vs. Shifted), whereas for the direct comparison between listening and production only non-shifted trials were used and Condition represented the modality (Listening vs. Produced).), Location (IN vs. OUT), Age, and PTA as covariates, with subject modelled as a random intercept.

For this stage of the study, we focused on demonstrating the feasibility of neural measurements under naturalistic conditions rather than performing detailed condi-tion-based comparisons. Consequently, the analysis of EEG responses was conducted using a sophisticated encoding approach (TRF analysis see above) to capture the temporal alignment between neural activity and continuous speech or gait signals. We deliberately refrained from applying complex statistical contrasts across conditions, as our primary goal was to assess whether reliable temporal response functions could be obtained during walking and natural conversation.

## Results

### Walking, listening, and speaking behaviour across tasks and environments

We analysed participants’ gait behaviour in terms of their walking velocity across four conditions: baseline (walking without any task), focused listening (listening to an audiobook while detecting repeats), the DTT (digit triplet test; listening to spo-ken digits in noise), and natural conversation (dialogue between the participant and the experimenter).

Participants walked faster in the baseline compared to the DTT (b = 0.07, SE = 0.013, t (180) = 5.8, p < .001) and the conversation (b = 0.09, SE = 0.013, t (180) = 7.7, p < .001). However, we found no significant differences between baseline and audiobook (b = - 0.02, SE = 0.013, t (180) = 1.4, p = 0.61). The pattern of results is visualized in Figure 2A.

**Figure 2.**
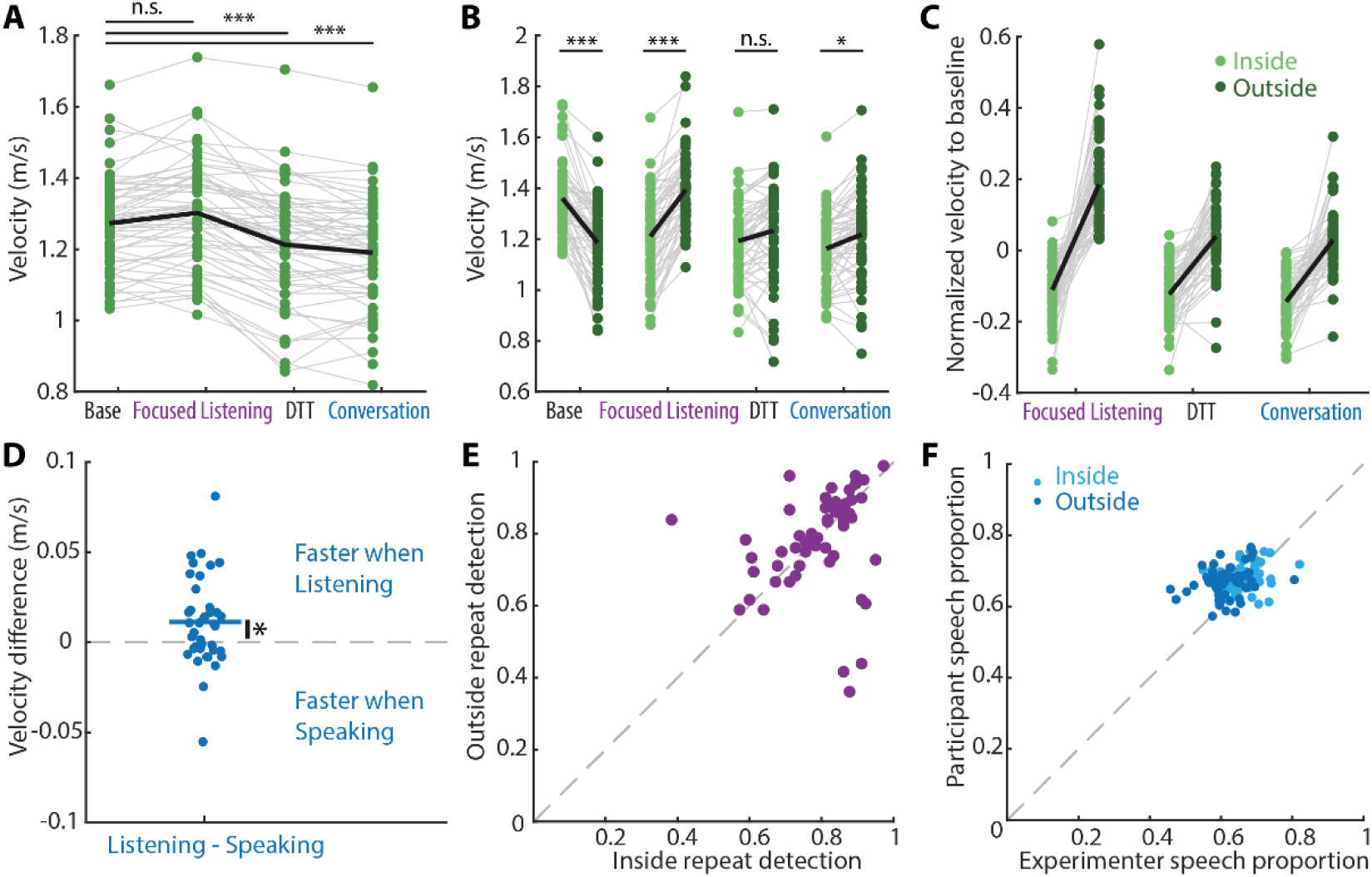
Context dependency of walking speed and behavioural perfor-mance. (A) Individual (green circles, gray lines) and average (thick black line) walking ve-locities across the four experimental conditions: baseline, focused listening, digit triplet test (DTT), and conversation. (B) Walking velocity separated by environ-ment (light green = inside, dark green = outside). Black lines show the mean per condition. (C) Velocity normalized to baseline, separated by environment. Each dot represents one participant; gray lines connect same participant. (D) Walking velocity differences between listening and speaking. Dots show indi-vidual participants; the solid blue line indicates the mean. (E) 45° plot of partici-pants’ focused listening (repeat detection) performance in indoor and outdoor en-vironments. (F) 45° plot of participant and experimenter speech proportions dur-ing the conversation task.

Participants performed the four walking tasks under two environmental conditions: indoors (hallway of the research building) and outdoors (courtyard in front of the building). We observed a strong interaction between task condition and environ-ment (F (420) = 73, p < .001). Given this interaction, we examined the effects in more detail (see Figure 2B).

Participants walked faster indoors compared to outdoors during the baseline con-dition (b = 0.20, SE = 0.019, t (420) = 10.7, p < .001). Interestingly, this pattern reversed for the auditory dual-task conditions. Participants walked slower indoors than outdoors during the audiobook task (b = –0.18, SE = 0.019, t (420) = –9.7, p < .001) and the conversation task (b = –0.06, SE = 0.019, t (420) = –3.2, p = .047). For the DTT, no significant difference between indoor and outdoor walking was observed (b = –0.04, SE = 0.019, t (420) = –2.1, p = .09).

In Figure 2C, the conditions are shown normalized to the baseline. Visual inspec-tion suggests that the differences between indoor and outdoor walking are further accentuated.

A closer examination of the conversation condition—specifically comparing peri-ods when participants were listening versus speaking (Figure 2D)—revealed a sig-nificant difference between the two states. Participants walked faster while listen-ing than while speaking (b = –0.01, SE = 0.005, t (151) = –2.1, p = .04).

Participants performed well in the repeat detection task during focused listening (mean inside = 0.79, 95% CI [0.76, 0.82]; mean outside = 0.78, 95% CI [0.74, 0.81]; Figure 2D), indicating that they were attending to the audiobook in this task.

During the conversation, participants and experimenters spoke roughly equal amounts of time (mean speech proportions with 95% CI: inside: participant: 0.686 [0.676, 0.696], experimenter: 0.663 [0.676, 0.696]; outside: participant: 0.667 [0.656, 0.679], experimenter: 0.612 [0.596, 0.628]). This indicates that the intended 50/50 split between speaking and listening was achieved, providing approximately equal amounts of data for participant (speaking) and experimenter (listening) in subsequent analyses. The proportions do not sum to 1 due to overlapping speech during the conversation.

### Speaking, not listening, drives speech–gait coupling

For the speech–gait coupling analysis, we used continuous representations of par-ticipants’ gait and speech signals. Figure 3A (upper panel) illustrates the continu-ous representation of a single participant’s gait, and the lower panel shows an ex-ample of the same participant’s speech envelope. Figure 3B displays the peak fre-quencies of gait and speech across all conditions and participants.

**Figure 3.**
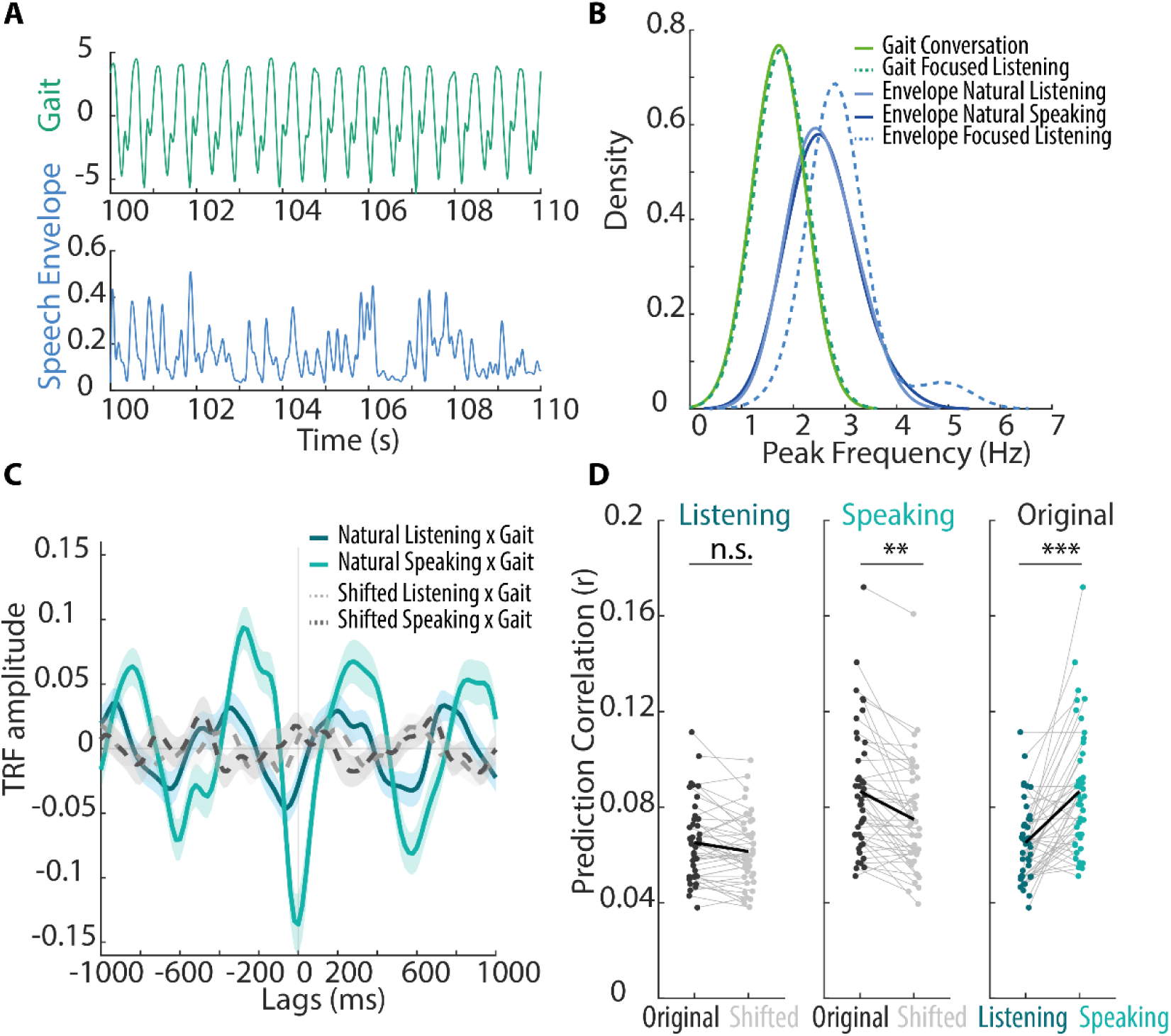
Speech–gait coupling emerges during speaking, not listening. (A) Example time series of gait (top, green) and the speech envelope (bottom, blue) from a natural listening segment. (B) Distributions of peak frequencies for gait (green) and speech envelope (blue) signals across conditions. Solid lines show natural conversation and natural speaking/listening, while dotted lines show focused listening. (C) Temporal response functions (TRFs) linking gait to speech during natural listening (dark blue) and natural speaking (cyan). Dotted lines indi-cate time-shifted control analyses. Shaded areas represent SEM across partici-pants. (D) Prediction correlations (r) for the listening (left) and speaking (right) conditions, comparing original data with time-shifted controls, and showing the comparison between original listening and speaking. Gray lines indicate individual participants; black lines represent group averages.

Figure 3C shows the TRFs for the coupling between the speech envelope and participants’ gait across all participants, separated for periods when participants were listening versus speaking. In addition, as a control, the TRFs for time-shifted speech envelope regressors are depicted. By visual inspection, the TRF for the speaking–gait coupling shows stronger components compared to the time-shifted control, particularly around time lags near zero, and the overall shape is more pronounced. In contrast, the TRF for the listening–gait coupling shows smaller components, also relative to the time-shifted control. Statistical analyses were performed on the prediction accuracies shown in Figure 3D.

Prediction accuracies for listening (original vs. time-shifted) and speaking (original vs. time-shifted) are shown in Figure 3D. Comparison of the listening condition revealed no significant difference between the original and time-shifted signals (b = –0.002, SE = 0.004, t (170) = –0.51, p = .5). In contrast, prediction accuracies were significantly higher for the original speaking signal compared to its time-shifted control (b = –0.013, SE = 0.004, t (170) = –2.79, p = .006). The coupling between speaking and gait was significantly stronger than the coupling between listening and gait (b = 0.027, SE = 0.006, t(170) = 4.66, p < .001). At first glance, the strong coupling between gait and speech could be attributed to a potential confound: that steps were also recorded by the microphones. However, this concern is mitigated by the use of speech separation algorithms, which detect speech and inherently exclude other noise sources. Moreover, if this were a significant confound, the listening condition should also exhibit similarly strong coupling, which it does not. Another possible confound is that the motion units may capture some chin movement. However, we extracted the gait signal from the eargait preprocessing software, which has already been filtered and pre-processed to isolate the relevant gait frequencies. Additionally, the analysis shown in Figure 3B confirms that the peak frequencies of gait and speech are distinct.

### Capturing listening-related neural responses while walking is feasible

We examined neural responses across different listening contexts. In the focused listening condition, participants listened to an audiobook while detecting repeated segments (Fig. 4A). This condition served as a baseline for neural responses during conversational listening. In the timing-uncorrected TRFs, we observed a positive peak at approximately 150–200 ms, followed by a negative peak around 200–300 ms. The single-subject TRF component map illustrates subject-specific TRF components, sorted by individual N1 latency. Following the realignment procedure described by Carta et al., we corrected for timing delays introduced by the technical recording setup (see inset in Fig. 4). Specifically, the individual N1 component of each participant was aligned to a time lag of 100 ms. As noted by Carta et al., this procedure tends to inflate the N1 component, and TRF latency values should not be interpreted quantitatively. After timing correction, the TRFs not only revealed a pronounced N1 component but, importantly, also exhibited the canonical auditory attention–related TRF morphology, including a P1–N1–P2 complex. Corresponding scalp topographies showed a fronto-central distribution, consistent with neural activity associated with attentive speech processing.

**Figure 4.**
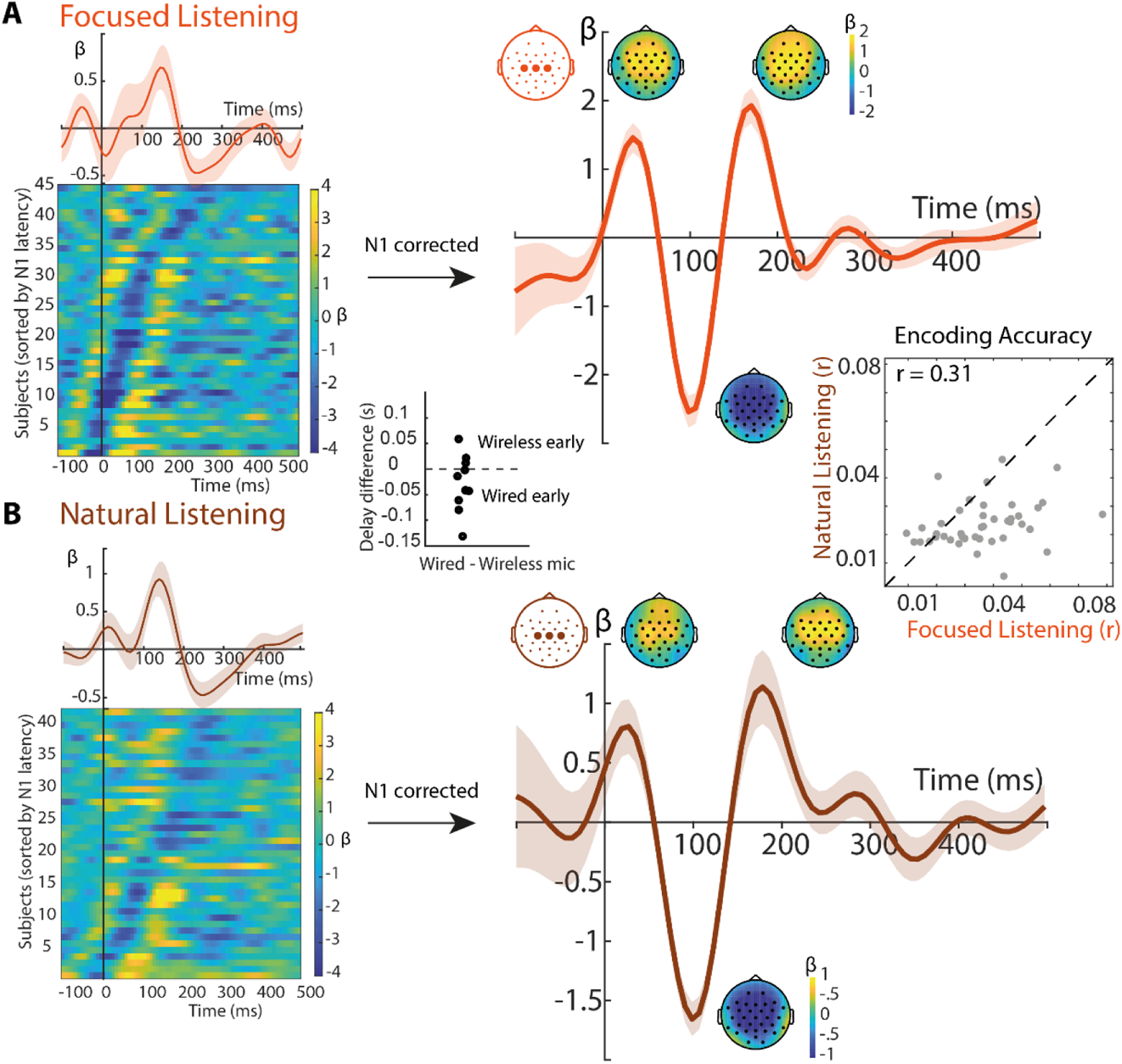
Temporal response function (TRF) during listening. (A) Focused listening. TRF β-weights for the focused listening condition, averaged across subjects and channels of interest (Cz, C3, and C4). Shaded areas indicate the standard error of the mean across subjects for each time lag. Scalp topographies show averaged TRF β-weights for selected time windows. On the left, subject-by-time β-weights are shown, sorted by individual N1 latency; N1-un-corrected, grand-averaged TRFs are displayed on the right, N1-corrected, grand-averaged TRFs are displayed. (B) Natural listening. TRF β-weights for the natural listening condition (conversation), averaged across subjects and channels of interest (Cz, C3, and C4). Shaded areas indicate the standard error of the mean across subjects for each time lag. Scalp topographies show averaged TRF β-weights for selected time windows. As in (A), subject-by-time β-weights sorted by N1 latency are shown on the left, and N1-corrected, grand-averaged TRFs are shown on the right. Insets. The left inset shows the measured delay difference between the wireless (used for stimulus recording) and wired microphones. The right inset shows encoding accuracy (correlation coefficient r) for electrodes Cz, C3, and C4, computed over time lags from −100 to 500 ms, comparing focused and natural listening conditions.

During natural listening (Fig. 4B), neural responses exhibited a pattern highly sim-ilar to that observed in the focused listening condition, even in the timing-uncor-rected TRFs. Specifically, we observed a positive peak at approximately 150–200 ms followed by a negative peak around 200–300 ms. After N1-based timing cor-rection, the TRFs again revealed the canonical P1–N1–P2 component structure. The corresponding scalp topographies closely resembled those observed during focused listening, showing the expected fronto-central distribution, albeit with gen-erally reduced magnitudes. Together, these results indicate that the TRF structure is largely preserved during natural, conversational listening, despite an overall at-tenuation of response strength compared to focused listening.

Encoding accuracy (Fig 4. Inset) was higher in focused listening (M = 0.036, 95% CI [0.031, 0.041], n = 42) compared to natural listening (M = 0.025, 95% CI [0.023, 0.028], n = 42). Furthermore, encoding accuracy showed a moderate positive correlation between the two conditions (r = 0.31), indicating that participants who exhibited larger encoding in one task tended to also have higher encoding in the other.

### Speaking- and gait-related TRFs and their encoding accuracy

We also examined the TRFs to natural speaking and gait, although primarily serv-ing as controls for potential movement artifacts for the listening conditions. The natural speaking TRF was time-shifted based on the N1 latency correction derived from the natural listening condition. During conversation in the natural speaking condition (Fig. 5A), we observed a pronounced positive peak around 30 ms, which was substantially larger in amplitude compared to the listening conditions. The scalp topography of this early peak was more pronounced over peripheral chan-nels, suggesting that it likely reflects a combination of predictive motor-related ac-tivity and residual artifacts from jaw movements, rather than purely neural pro-cesses. The inset in Fig. 5A shows the uncorrected natural speaking TRF.

**Figure 5.**
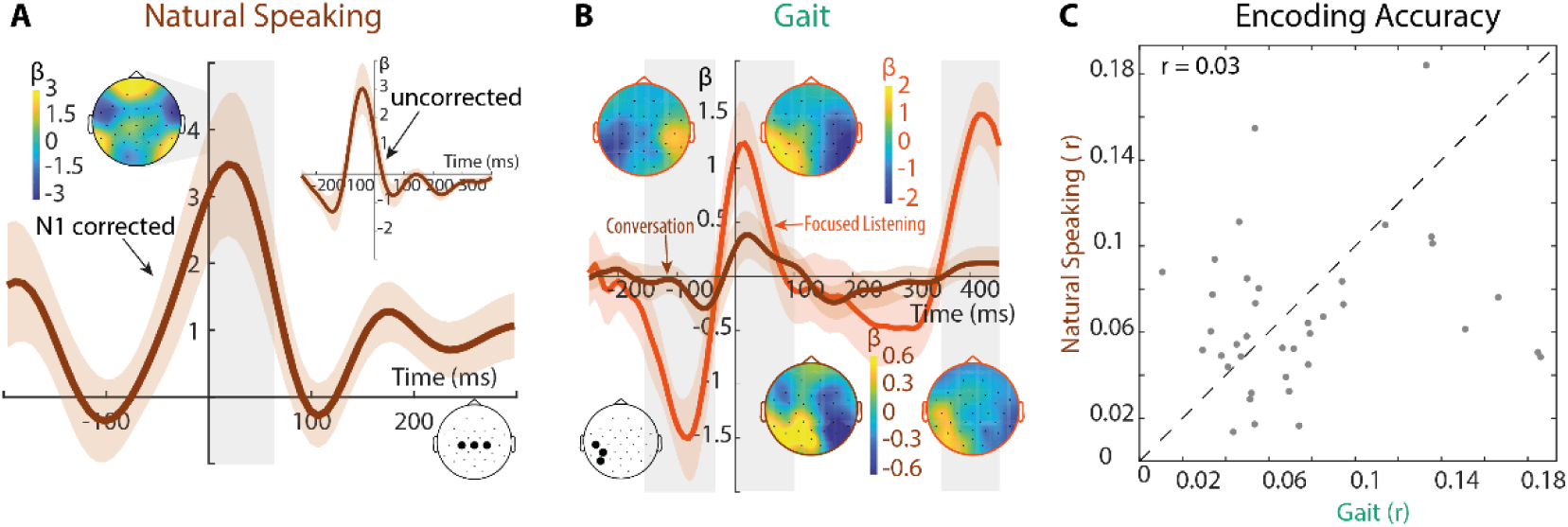
Temporal response function (TRF) to natural speaking and gait (A) Natural speaking. TRF β-weights for N1-corrected natural speaking, averaged across subjects and channels of interest (Cz, C3, and C4). Shaded areas indicate the standard error of the mean across subjects for each time lag. Scalp topographies show averaged TRF β-weights for selected time windows. Inset shows un-corrected TRF. (B) TRF to gait during the conversation and focused listening. TRF β-weights are aver-aged across subjects and channels of interest (T7, P7 and CP5). Shaded areas depict the standard error for each time lag across subjects. Topographies show averaged TRF β-weights for highlighted time windows. (C) Encoding accuracy. Correlation coefficient r for all electrodes, computed over time lags from −200 to 400 ms, comparing natural speaking and gait during the conver-sation task.

The gait TRF, measured while participants were simultaneously performing the fo-cused listening or natural conversation task (Fig. 5B), revealed dynamics distinct from the auditory TRFs, with a negative peak around –80 ms and positive peaks near 30 ms, reflecting neural activity related to gait. The topographic distribution was lateralized, likely reflecting left- and right-step activity. While the time course and topography were generally consistent across tasks, including focused listening and natural conversation, peak magnitudes were larger in the focused listening task.

Encoding accuracy differed between the gait and speaking conditions. The gait condition showed a mean encoding accuracy of 0.086 (95% CI [0.066, 0.105], n = 42), whereas the speaking condition exhibited a lower mean encoding accuracy of 0.066 (95% CI [0.055, 0.077], n = 42). Overall, encoding accuracy values for both gait and speaking were higher than those observed for the listening conditions, potentially reflecting contributions from movement-related signals or residual arti-facts.

## Discussion

In the present study, we aimed to investigate how neural, behavioural, and motor processes interact during natural communication and movement. Using simultane-ous EEG, audio, and motion recordings, we examined how listening, speaking, and walking are coordinated in real-world conditions. Our primary goal was to assess whether reliable neural responses to natural conversational speech can be meas-ured while participants are in motion. We further explored how environmental, and task demands influence gait behaviour, and whether rhythmic coupling emerges between speech and gait.

### Gait slowing in cognitively demanding auditory tasks

A first central finding is that participants’ gait behaviour varied robustly across the four walking tasks. Walking speed decreased when participants engaged in cogni-tively demanding auditory tasks, particularly during the digit triplet test and during natural conversation. This aligns with dual-task literature showing that speech per-ception under challenging conditions draws on cognitive resources that are also needed for gait control (Li et al., 2014; Verghese et al., 2007; Yogev-Seligmann et al., 2008). In contrast, listening to an audiobook while performing a simple repeat-detection task did not significantly slow participants down relative to baseline walk-ing. This suggests that not all forms of listening impose the same cognitive load: continuous speech can be processed while walking with minimal interference, whereas tasks that require active speech formulation, as in conversation, impose additional cognitive demands that lead to measurable motor consequences.

### Indoor–Outdoor differences in gait during auditory tasks

An equally important result is the strong interaction between participants gait ve-locity and environmental context. Indoors, baseline walking was faster than out-doors, likely reflecting a more predictable and constrained environment (smooth floor, straight corridor) that allows participants to adopt a more efficient gait pattern. Interestingly, this pattern reversed during the auditory dual-task conditions: out-doors, participants walked faster than indoors while listening to the audiobook and during conversation. A plausible explanation for this reversal relates to differences in room acoustics between the two environments. The indoor corridor was highly reverberant, whereas the outdoor courtyard approximated a free-field condition with minimal reflections. Increased reverberation is known to degrade the temporal structure of speech and introduce delayed energy that can resemble repetitions or echoes (Kjellberg, 2004; Moore, 2007; Reinhart & Souza, 2016; Xia et al., 2018). Such acoustic smearing likely made the auditory tasks—especially focused listen-ing to the audiobook—more cognitively demanding indoors than outdoors.

Under these conditions, participants may have allocated more attentional re-sources to speech processing indoors, leaving fewer resources available for gait control, which in turn resulted in slower walking. In contrast, the outdoor free-field environment presented a clearer acoustic signal and thereby allowing participants to maintain higher walking speeds. This interpretation also aligns with the particu-larly strong indoor–outdoor difference observed in the focused listening task, where reverberation is most likely to interfere with the detection of repeated phrases or temporally shifted acoustic events.

Together, these results highlight that neither cognitive nor environmental con-straints alone explain flexible gait modulation; rather, it is their interaction that shapes behaviour in naturalistic contexts.

### Differential gait demands of speaking and natural listening

Within the natural conversation, we observed a modulation of gait depending on the communicative role: participants walked faster while listening than while speak-ing. While previous research has mostly examined scripted or laboratory-based speech tasks, our results suggest that, even in spontaneous face-to-face conver-sations with turn-taking, speech production may place greater demands on cogni-tive and motor resources than listening, thereby affecting gait (Atakanova et al., 2023; Beurskens et al., 2016; Helfer et al., 2023; Holtzer et al., 2014; Li et al., 2014). Formulating an utterance in real time involves syntactic planning, word re-trieval, articulatory control, and breathing coordination — all of which compete with the resources needed to maintain stable locomotion (Houde & Jordan, 1998; Inde-frey & Levelt, 2004; Macneilage, 1980). The roughly balanced speaking and natu-ral listening proportions between participant and experimenter confirm that the con-versational structure worked as intended. Overall, these findings extend prior dual-task research by demonstrating that the asymmetry in motor costs between pro-duction and natural listening also emerges in highly ecological, interactive settings.

### Differential coupling of gait with speaking versus natural listening

The present results provide converging evidence for a systematic coupling between participants’ gait and their speech production, while revealing comparatively weak or absent coupling during natural listening. Using continuous representations of gait and speech, we observed clear temporal response functions (TRFs) linking the speech envelope to gait dynamics during speaking, with components that ex-ceeded those obtained from time-shifted control regressors. This pattern suggests that the observed coupling is a genuine temporal alignment between motor pro-cesses involved in speech production and locomotion.

In contrast, during natural listening periods, TRFs showed only small components, and prediction correlation did not differ from the time-shifted control (unreliable time-lags). This dissociation between speaking and natural listening indicates that the interaction between speech and gait is driven primarily by active speech pro-duction rather than by listening to the conversational partner. One interpretation is that speech production recruits motor and respiratory rhythms that become tem-porally aligned with the rhythmic structure of gait. In addition to these systems, cardiac rhythms may also play a role: heart rate oscillates within a low-frequency range that overlaps with both respiratory and locomotor rhythms, and cardiorespir-atory–locomotor coupling has been documented in several motor domains (Abbasi et al., 2023; Di Pasquasio et al., 2025; Juvin et al., 2022; Kirby et al., 1989). These interacting rhythms—respiratory, articulatory, and cardiac—may jointly act as driv-ers of the observed speech–gait coupling, consistent with broader principles of rhythmic motor coordination across multiple physiological systems.

Future studies will be needed to test this possibility more directly. Simultaneous recordings of respiration, heart rate, articulatory movements, and gait would allow disentangling their relative contributions to the coupling. Furthermore, experi-mental manipulations that modulate respiratory or cardiac rhythms (e.g., paced breathing, induced changes in heart rate) could provide causal evidence on how these physiological oscillators shape the interaction between speech production and locomotion.

An alternative interpretation—that gait–speech coupling is driven by acoustic con-tamination of the speech recordings—is unlikely for several reasons. First, speech separation algorithms were applied to isolate the speech envelope, substantially reducing the contribution of footstep-related vibrations or environmental noise. If step-related artifacts were still present, one would expect to find similarly strong coupling during listening segments, where no articulation occurs and only external auditory input is recorded by the microphones. This was not the case: the natural listening condition showed no significant coupling. Second, gait signals were derived from the eargait preprocessing pipeline, which includes filtering and feature extraction designed specifically to isolate gait-related oscillations. Thus, the gait regressor reflects biomechanical movement rather than acoustic contamination or local vibrations.

A further potential confound is that the motion units might capture jaw movements during speech. While we cannot entirely exclude this possibility, several aspects of the data argue against it being the driving factor. The extracted gait signal is dom-inated by frequencies characteristic of locomotion, and analysis of peak frequen-cies demonstrated clear separability between gait and speech rhythms. If jaw movements were substantially contaminating the motion sensors, these frequen-cies would likely overlap or show stronger correspondence across both speaking and listening conditions. Instead, the observed coupling is specific to speaking and aligns with the temporal structure of speech production rather than with the static or limited jaw motion present during natural listening.

Taken together, these findings point to a functional coordination of speech and gait during speech production. The stronger coupling during speaking suggests that speech motor processes interact with whole-body locomotor rhythms, potentially through shared neural oscillators or biomechanical constraints that favor rhythmic alignment. The absence of comparable effects during natural listening suggests that auditory processing alone does not impose rhythmic structure on gait. These results could have implications for understanding naturalistic communication dur-ing walking and for investigating motor–speech interactions in populations with gait or speech impairments.

### Feasibility of neural speech tracking during walking and natural conversa-tion

The central finding of this study is that reliable neural responses to speech enve-lopes can be obtained while participants are walking, both during focused listening and during natural conversational listening. Despite the increased motor activity and reduced experimental control compared to laboratory paradigms, the canoni-cal P1–N1–P2 TRF morphology was clearly observable in both conditions after timing correction. The corresponding scalp topographies showed the expected fronto-central distribution associated with auditory cortical processing and atten-tional engagement, consistent with prior work conducted in stationary laboratory environments (Brodbeck et al., 2018; Crosse et al., 2016; Di Liberto et al., 2015; Ding & Simon, 2012; Orf et al., 2023; Zion Golumbic et al., 2013).

Importantly, these results extend previous real-world EEG studies that primarily relied on passive listening to audiobooks or controlled speech streams (e.g., Straetmans et al., 2024) by demonstrating that neural speech tracking is preserved during genuine conversational interactions. Natural conversation differs fundamen-tally from audiobook listening in that it involves dynamic turn-taking, spontaneous speech production, variable prosody, and rapid alternations between listening and speaking. The presence of robust listening-related TRFs under these conditions suggests that neural tracking of speech envelopes generalizes beyond highly con-trolled listening paradigms and remains measurable even during highly dynamic, socially interactive behaviours, which provides a promising basis for future work explicitly targeting these conversational dynamics.

While encoding accuracy was reduced during natural listening compared to fo-cused listening, this attenuation likely reflects the increased cognitive and percep-tual complexity of conversation, including divided attention, unpredictable speech dynamics, and concurrent motor demands. In addition, speech signals during nat-ural conversation were obtained using a deep neural network–based speech sep-aration approach (Ravanelli et al., 2021). Although this method enables the isola-tion of individual speakers in realistic acoustic environments, it is not free of resid-ual distortions and separation errors. Such imperfections may further reduce the fidelity of the extracted speech envelope and thereby contribute to lower encoding accuracy. Nevertheless, the moderate correlation in encoding accuracy across conditions indicates stable individual differences in neural speech tracking ability, supporting the robustness of the approach.

### Speaking- and gait-related TRFs: artifacts, neural contributions, and unified modelling

In addition to listening-related TRFs, we estimated TRFs for natural speaking and gait to characterize neural and non-neural signals associated with speech produc-tion and movement that could otherwise confound the interpretation of listening-related responses. These analyses primarily served a control function within our encoding framework.

Speaking-related TRFs were dominated by an early, large-amplitude positive peak around 30 ms with a predominantly peripheral scalp distribution. This pattern is unlikely to reflect purely auditory cortical processing and likely arises from a com-bination of motor-related activity, bone-conducted self-generated speech, and re-sidual jaw and facial muscle artifacts.

Gait-related TRFs exhibited distinct temporal dynamics and lateralized scalp to-pographies consistent with step-related activity. Importantly, the overall TRF time course and topographic distribution were qualitatively similar to gait-related neural responses reported by (Herbert & Munz, 2020), suggesting that the observed ef-fects may, at least in part, reflect genuine neural contributions to gait control. A key methodological difference, however, is that we modelled TRFs based on the con-tinuous gait signal, which naturally includes both left and right initial contacts. This continuous approach likely explains the observed lateralization in scalp topogra-phies, in contrast to Herbert and Munz, who examined gait-related event-related potentials time-locked exclusively to left initial contacts.

Encoding accuracy for speaking and gait exceeded that observed for listening. Ra-ther than indicating stronger cortical encoding, these elevated values likely reflect the high signal-to-noise ratio of movement-related signals and residual artifacts that are tightly time-locked to the respective regressors. While the similarity to pre-viously reported gait-related ERPs supports the presence of genuine neural com-ponents in the gait TRFs, we cannot exclude contributions from movement-related artifacts. Disentangling neural and non-neural sources of gait-related TRFs will re-quire further studies using complementary approaches.

A key methodological contribution of this study is the use of a unified encoding framework in which listening, speaking, and gait-related signals are modelled jointly. By explicitly accounting for variance driven by movement and speech pro-duction, listening-related TRFs can be interpreted more confidently without relying on aggressive artifact rejection. This approach is particularly important for conver-sational paradigms, where listening and speaking are tightly interwoven, and demonstrates that meaningful neural speech tracking can be obtained even under naturalistic, real-world conditions.

## Limitations of the study

While the present study demonstrates the feasibility of measuring neural speech tracking and gait-related responses under naturalistic, real-world conditions, sev-eral limitations should be noted. First, the uncontrolled nature of outdoor and semi-naturalistic environments introduces variability that is difficult to fully account for, in contrast to highly controlled laboratory paradigms. Factors such as fluctuating background noise, variable walking surfaces, and spontaneous environmental events may contribute to additional cognitive and perceptual demands, potentially affecting both gait and neural responses.

Second, the current setup did not include complementary physiological and behav-ioural measurements that could have enhanced our understanding of the observed effects. For example, simultaneous recordings of respiration, heart rate, eye-track-ing, or full-body kinematics could help disentangle contributions from autonomic rhythms, articulatory processes, and motor coordination.

Third, the technical complexity of the mobile recording setup introduced small but non-negligible delays in stimulus presentation and EEG recording, limiting the in-terpretability of absolute TRF latencies. Due to the technical delays introduced by the complex mobile setup, absolute TRF latencies cannot be interpreted, and tim-ing corrections only allow for qualitative assessment of the canonical components.

Finally, these challenges underscore the importance of robust and well-synchro-nized technical solutions when conducting EEG and behavioural research “in the wild”. Despite these limitations, the present findings highlight that meaningful neu-ral and behavioural signals can still be extracted under real-world conditions, in-cluding during natural conversational interactions while walking.

## Conclusion

Overall, despite the challenges inherent to real-world recordings, our study shows that meaningful neural tracking of speech can be captured during natural conver-sations while participants are moving. These results demonstrate that the brain robustly aligns with the temporal structure of speech even in complex, dynamic environments. By bridging laboratory findings with ecologically valid settings, this work provides a foundation for future investigations of auditory processing and cognition in everyday life.

## Acknowledgements

Research was supported by a research grant from ORCA Labs Europe, Erlangen, Germany, to JO.

